# Subject identification using edge-centric functional connectivity

**DOI:** 10.1101/2020.09.13.291898

**Authors:** Youngheun Jo, Joshua Faskowitz, Farnaz Zamani Esfahlani, Olaf Sporns, Richard F. Betzel

## Abstract

Group-level studies do not capture individual differences in network organization, an important prerequisite for understanding neural substrates shaping behavior and for developing interventions in clinical conditions. Recent studies have employed “fingerprinting” analyses on functional connectivity to identify subjects’ idiosyncratic features. Here, we develop a complementary approach based on an edge-centric model of functional connectivity, which focuses on the co-fluctuations of edges. We first show whole-brain edge functional connectivity (eFC) to be a robust substrate that improves identifiability over nodal FC (nFC) across different datasets and parcellations. Next, we characterized subjects’ identifiability at different spatial scales, from single nodes to the level of functional systems and clusters using k-means clustering. Across spatial scales, we find that heteromodal brain regions exhibit consistently greater identifiability than unimodal, sensorimotor, and limbic regions. Lastly, we show that identifiability can be further improved by reconstructing eFC using specific subsets of its principal components. In summary, our results highlight the utility of the edge-centric network model for capturing meaningful subject-specific features and sets the stage for future investigations into individual differences using edge-centric models.

## INTRODUCTION

Over the past several decades, the field of neuroimaging has leveraged powerful computational methods to develop standardized, population-level descriptions of brain morphology and functional organization [1, 2]. These approaches have facilitated group-level and cross-sectional comparisons of brains [3, 4], enhancing our knowledge of the functional and neuroanatomical underpinnings of cognition [5, 6], brain development [7, 8], and neuropsychiatric disease [9]. However, these efforts have emphasized group-level effects at the expense of individual brains, whose organization is personalized and idiosyncratic [10–12].

Recently, several important studies have begun to shift focus away from group-level analyses and onto single subjects [13, 14]. The aim of this line of research is to build comprehensive maps of individuals’ brains [11, 12, 15, 16], with the hope that inclusion of personalized details will add clarity to brain organization, brain-behavior relationships [17–21], and inform treatment of neuropsychiatric disorders by helping design more efficacious and targeted interventions [22, 23].

One particular strand of this research focuses on mapping the features of brain networks that are idiosyncratic to individuals. Like fingerprints, these features are capable of distinguishing one person’s brain from that of another [14, 24]. Brain network fingerprints serve as reliable substrates of individuals [14, 25] that are stable over time [26–28], across subsets of FC data [29], and across acquisition sites [30]. In addition, identifiable characteristics have shown clinical diagnostic potential [31] and have proven useful for the classification of individual behaviors and cognitive states [32, 33].

To date, most fingerprinting analyses have focused on features derived from brain networks in which nodes represent positioned electrodes [34] or brain regions [14, 35]. Recently, we proposed an alternative model of brain connectivity that focuses on interactivity among a network’s connections (or “edges”) [36–39]. We refer to these patterns as edge functional connectivity (eFC). Adopting an edge-centric perspective has been fruitful in other scientific disciplines, e.g. in uncovering the overlapping community structure of complex biological and social networks [40, 41]. Similarly, eFC has provided a new window into studying the organization of brains, including the overlapping community structure [36] and how participation of functional systems vary across communities [38]. However, it remains unclear how eFC compares with traditional nodal functional connectivity (nFC) in terms of its ability to convey individual-specific information.

Here, we extend connectome-based fingerprinting to edge-centric networks. Using functional imaging data from two independently acquired datasets (the Midnight Scan Club[15] and the Human Connectome Project [42]; MSC and HCP, respectively), we compare the performance of whole-brain nFC and eFC on subject identification, demonstrating that with sufficient amounts of data, eFC enables greater and more robust identifiability than nFC. Next, we investigated the system- and node-level drivers of the improved identifiability in eFC, focusing on system-specificity using a “leave-one-node-out” approach. We found system nodes and edges associated with heteromodal brain systems to be the primary drivers of subject identification. Finally, we tested whether it was possible to optimize identifiability by reconstructing eFC and nFC using a restricted set of principal components. We found that reconstructed eFC significantly outperformed that of reconstructed nFC in terms of its optimized identifiability. Our work sets the stage for future studies to use eFC to develop network-based biomarkers for tracking inter-individual differences in behavior, development, and disease diagnosis.

## RESULTS

In this report, we systematically evaluate eFC and nFC in terms of differential identifiability and discuss their similarities and differences at regional (node) and subsystems (group of nodes) levels. Throughout this section, we analyze data from two high-quality independently acquired datasets: the Midnight Scan Club (MSC; [15, 28]), which consists of ten participants scanned ten times each, and 100 unrelated subjects scanned two times each from the Human Connectome Project (HCP; [42]).

### Identifiability using edge functional connectivity

Subject identification can be quantified using the measure “differential identifiability”, or *I*_*diff*_, which is calculated as the mean within-subject similarity minus the mean between-subject similarity of connectivity matrices [35]. Existing subject identification applications have relied on connectivity patterns derived from nFC and thus the idiosyncratic characteristics of eFC remains unknown. In this section we compare the identifiability of cortex-wide nFC and eFC and its dependence on the amount of data available.

First, we compared cortex-wide eFC and nFC in terms of subject identifiability. Briefly, this entailed estimating nFC and eFC separately for each of the 100 resting-state fMRI scans (10 subjects; 10 scans each) in the Midnight Scan Club dataset, and generating similarity matrices for each connectivity modality as the Pearson correlation between the upper triangle elements of subjects’ nFC or eFC matrices (Fig. 1*e*). We then estimated differential identifiability from these similarity matrices (Fig. 1*f*).

**FIG. 1.**
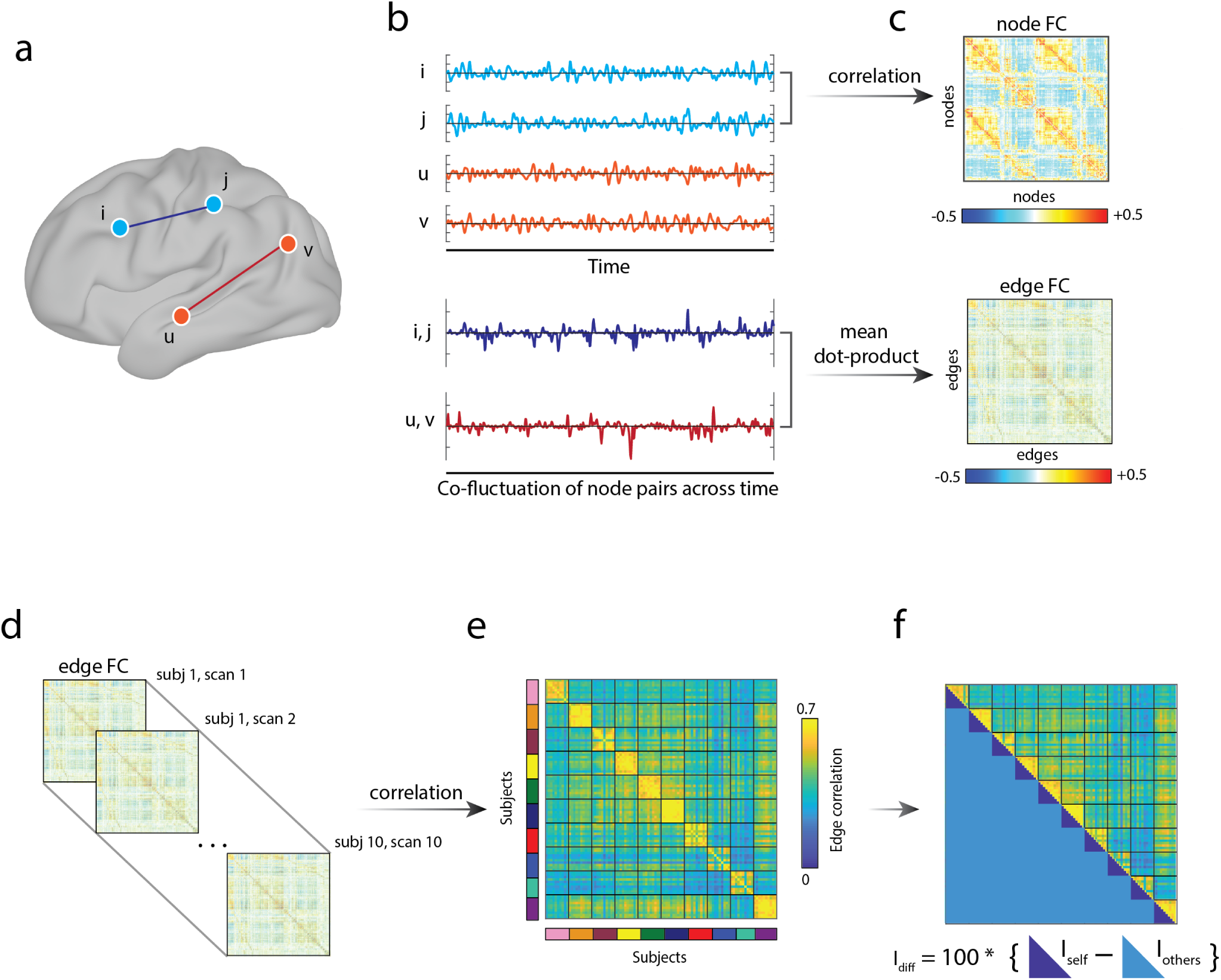
Schematic of differential identifiability in eFC. All panels from this figure were generated using data from the Midnight Scan Club dataset. (*a*) To illustrate the calculation of node and edge FC (nFC and eFC, respectively), consider four nodes: *i, j, u*, and *v*. nFC is defined as the pairwise correlation of regional activity. For nodes *i* and *j*, nFC is calculated by first z-scoring each nodes’ time series, computing the element-wise product, and averaging these values (panels (*b*) and (*c*); *top*). The same operation could be carried out for nodes *u* and *v*. eFC is calculated by first generating co-fluctuation time series for pairs of nodes. This involves computing their element-wise product, but omitting the averaging step (panel (*b*); *bottom*). Each co-fluctuation time series is defined for a pair of nodes. eFC is calculated as the temporal similarity (e.g. correlation, cosine similarity, etc.) of pairs of co-fluctuation time series ((*c*); *bottom*).(*d*) To calculate differential identifiability, (*e*) we extract the upper triangle elements of subjects’ eFC matrices and compute the spatial correlation of those elements, resulting in subject-by-subject similarity matrix. (*f*) Differential identifiability, *I*_*diff*_ is calculated as the mean within-subject similarity minus the mean between-subject similarity.

We found that eFC outperformed nFC, yielding greater values of *I*_*diff*_ when using whole brain derived functional networks (Fig. 2*a*). Similar results were also found when using different parcellations and in an independent dataset (Fig. S1). To visualize the within- and between-subject similarity, we used multi-dimensional scaling to project subjects and their scans into a two-dimensional space that approximately preserves the pairwise distance relationships encoded in the nFC and eFC similarity matrices (Fig. 2*b-c*). We note that these plots are for the purposes of visualization, only.

**FIG. 2.**
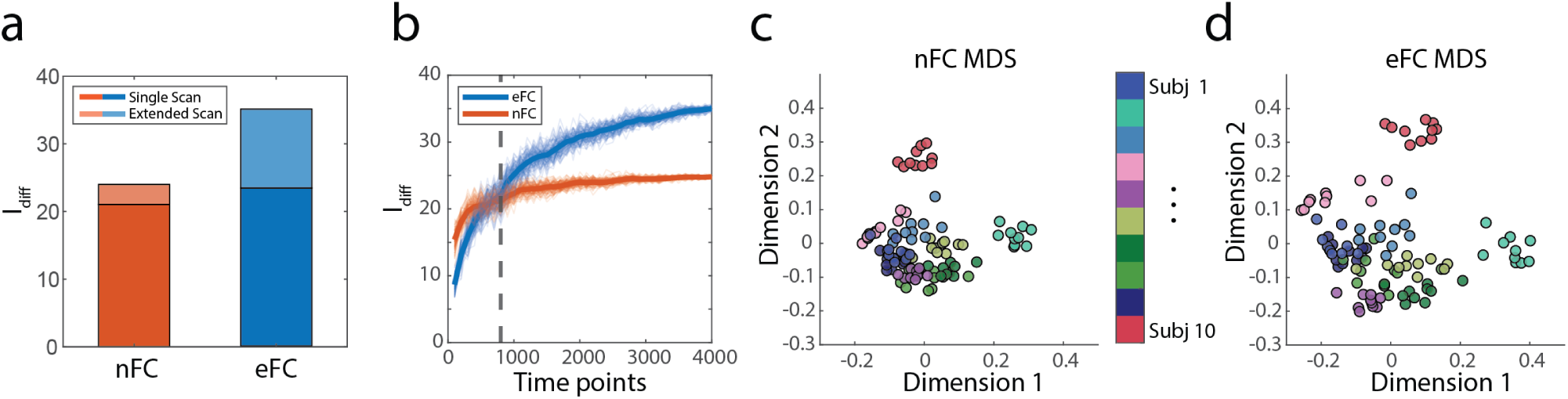
Subject identification of eFC and nFC and effect of scan length. All panels from this figure were generated using data from the Midnight Scan Club dataset with 200 node Schaefer parcellations. (*a*) *I*_*diff*_ of nFC and eFC for single scans (darker color) and for maximally concatenated scans (lighter color). (*b*) *I*_*diff*_ of eFC and nFC by scan length. The black dotted line (800 time points) indicates the length of the time series in which eFC significantly outperforms nFC. Thick blue and orange lines are the average *I*_*diff*_ s for 100 iterations and thin lines indicate each iteration’s *I*_*diff*_ by scan length. (*c*) Subjects’ scans plotted using multi-dimensional scaling for nFC. (*d*) Subjects’ scans plotted using multi-dimensional scaling for eFC. Each subject’s scan corresponds to a color on the colorbar between panels *c-d*.

In the previous analysis, we calculated eFC and nFC using approximately 30 minutes of data (the duration of scan session in the MSC dataset). Next, we tested whether subject identifiability was modulated by scan length, i.e. whether the value of *I*_*diff*_ varied as a function of the amount of data available [30, 43]. To test this, we created shorter or longer “sessions” by either dividing the existing runs into shorter, contiguous segments, or by concatenating data from multiple scans to form longer sessions. We varied the duration of artificial scan sessions in increments of 100 samples, starting with 100 and ending with 4000. This entire sampling procedure was repeated 100 times. We found that with fewer than 500 time points (approximately 20 minutes) *I*_*diff*_ was greater for nFC than eFC (*p <* 10^−6^, *t − test*); Fig. 2*a*. However, at 800 time points (approximately 30 minutes), eFC began to significantly outperform nFC (*p <* 10^−4^; *t − test*); Fig. 2*b*. We report similar results using different parcellations and datasets (Fig. S1).

Our results using eFC are in line with previous research where identifiability was increased with extended scan length using conventional nFC [30, 35]. Collectively, our findings indicate that, given sufficient amounts of data (approximately 30 minutes), eFC enables a more robust identification of subjects across sessions than nFC.

### Regional drivers of cortex-wide eFC identifiability

In the previous section, we found that eFC improved cortex-wide identifiability over nFC given a sufficient number of samples. In this section, we wanted to pin-point the brain regions that contributed to this improvement. To do this, we use a “leave-one-node-out” method to measure the relative impact that each brain region had on subject identification. We then summarized these results by grouping nodes according to canonical brain systems and assessing, statistically, the contribution of each system to the overall identifiability.

In order to trace identifiability in eFC back to brain regions, we first calculated the difference in *I*_*diff*_ estimated using an intact brain (all 200 nodes) with the *I*_*diff*_ estimated after iteratively removing individual nodes, one at a time. We also performed a similar analysis using edges instead of nodes, the results of which are reported in Fig. S2. We found that, when removed, cortical areas located in the control and temporoparietal network regions yielded decreased differential identifiability while removal of regions associated with somatomotor, limbic, and visual networks increased identifiability (Fig. 3*a*; note that for the sake of visualization, we invert the sign of Δ*I*_*diff*_). All nodes were also grouped according to their systems visualizing each node’s contribution to subject identifiability (Fig. 3*b*). Each node’s contribution was measured as the difference in *I*_*diff*_ without a particular node and subtracting the *I*_*diff*_ calculated from all 200 nodes. For example, in Fig. 3*b*, removing a single node from the control A system (represented as a single yellow dot) prior to eFC construction would result in a reduction in *I*_*diff*_, which we plot here as having a “positive” effect on *I*_*diff*_. In contrast, removing a single node from the limbic system yields an increase in *I*_*diff*_, hence we plot as a “negative” effect on *I*_*diff*_.

**FIG. 3.**
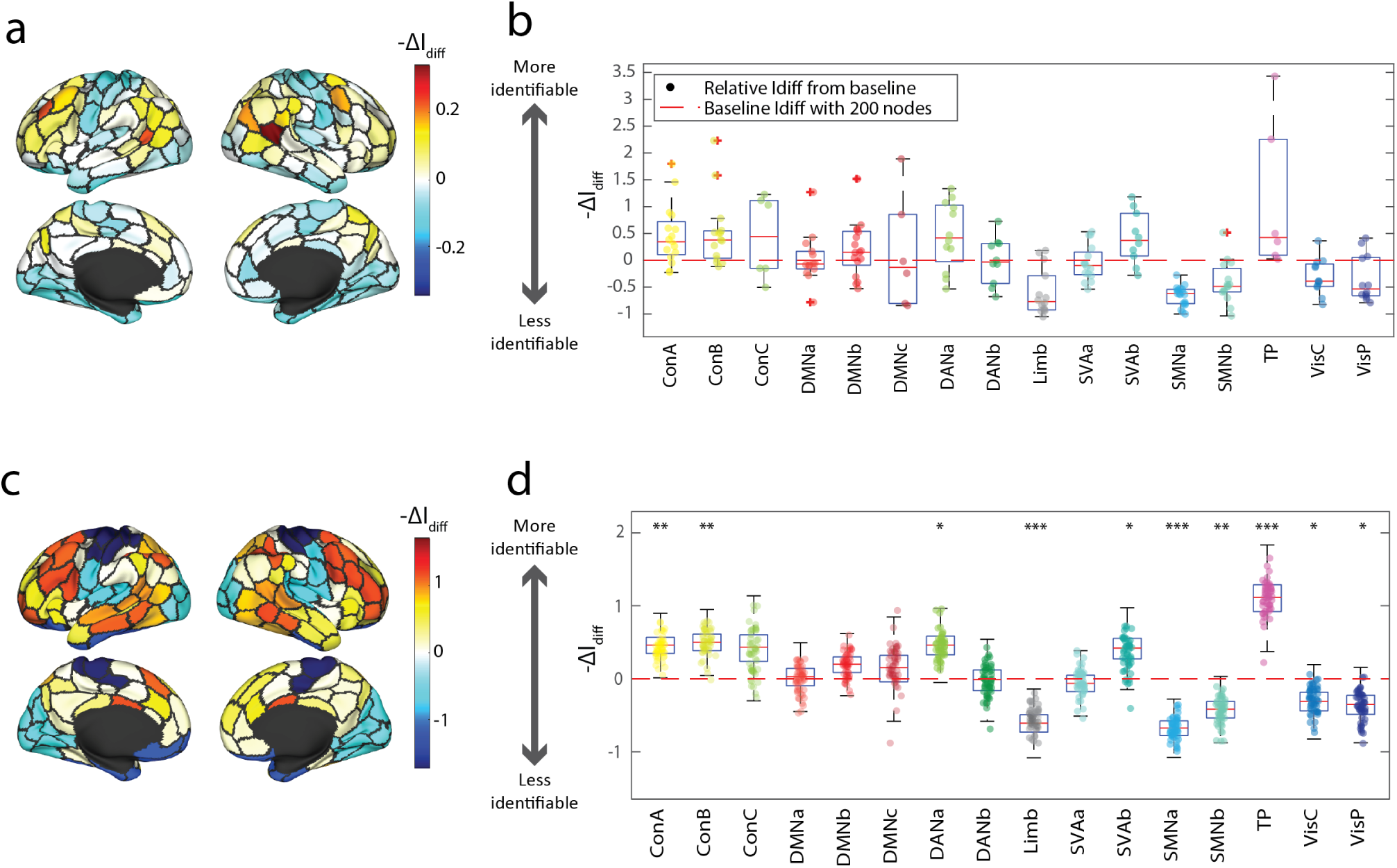
Nodal contribution to identifiability in eFC. (*a*) To assess regional contributions to *I*_*diff*_, we calculated the change in *I*_*diff*_ after removing all edges incident upon each of the *N* = 200 nodes. Here, we show the change in *I*_*diff*_ projected onto the cortical surface. (*b*) We plotted the relative impact of each node’s removal prior to eFC calculation on *I*_*diff*_ by brain systems. (*c*) Excluding entire system’s nodes prior to eFC construction reveals that specific systems contribute positively or negatively to *I*_*diff*_ and that (*d*) compared to random removal of matching numbers of nodes, have significantly different results on *I*_*diff*_ (Bonferroni-corrected *p*-values; * = *p <* 0.003, ** = *p <* 0.0006, *** = *p <* 0.00006). For visualization we inverted the Δ*I*_*diff*_ to *−* Δ*I*_*diff*_, which quantifies the relative effect of a node’s removal on the overall *I*_*diff*_. If a removal of a node results in reduced *I*_*diff*_, this node’s contribution to identifiability would be considered as “increasing *I*_*diff*_ ”.

Next, we tested which system’s nodes significantly influenced identifiability compared to random removal of matching numbers of nodes (Fig. 3*c – d*). Nodes were randomly reassigned to systems by randomly permuting system labels (10,000 iterations). We found that when removing nodes from the control A, control B, dorsal attention A, salience ventral network B, and temporoparietal networks significantly decrease identifiability than when randomly removing matching numbers of nodes (thus, the relatively positive effect of these systems in panel Fig. 3*d*). Also, we found that excluding nodes from limbic, somatomotor A and B, and central and peripheral visual networks significantly increase identifiability compared to randomly removing matching numbers of nodes (thus, the relative negative effect of these networks to *I*_*diff*_ in panel Fig. 3*d*).

In summary, we used a *leave-one-node-out* approach to uncover the regional drivers of whole-brain identifiability. We found that the inclusion of frontoparietal and superior temporal regions help increase identifiability, while somatomotor, limbic, and visual regions lead to reductions in identifiability. These results on system-level identifiability largely agree with prior research using conventional, node-centric functional connectivity [14, 44, 45], localizing idiosyncrasies of brain network organization to a specific subset of systems.

### Identifiability of systems and clusters in eFC

In the previous sections, we demonstrated that given an fMRI scan of sufficient duration eFC outperforms nFC in subject identification and that heteromodal brain regions compared to unimodal, contribute to higher identifiability. Here, we continued our investigation into the drivers of cortex-wide identifiability, focusing on the contributions of each functional brain system in eFC to identifiability. In this section, we aim to answer the questions: How do edges from single systems contribute to identifiability? How do clusters of edges perform in identifiability?

To address these questions, we first estimated *I*_*diff*_ using only connections associated with specific brain systems. In the case of nFC, this means calculating identifiability using only edges whose stub nodes are assigned to the same brain system [46]. We performed a similar operation using eFC using edge pairs subjected to the requirement that all four nodes associated with the two edges that comprise an eFC entry were assigned to the same system. In general, we found that system-specific *I*_*diff*_ for eFC and nFC was highly correlated (*R* = 0.9578, *p <* 10^−8^); Fig. 4*b*). We found that the greatest levels of *I*_*diff*_ were associated with temporoparietal, control A, and dorsal attention networks (Fig. 4*a*). In contrast, edge pairs in eFC that are solely from somatomotor B and default mode B networks were found to have lower *I*_*diff*_ (see Fig. S3 for system-level similarity matrices). We note that these single-system edge pairs represent only a small fraction (*<* 1%) of the eFC matrix and that the eFC incorporates a broader repertoire of edge pairs, some of which involve nodes originating in four distinct systems. Subject similarity matrices for single systems are included in Fig. S2.

**FIG. 4.**
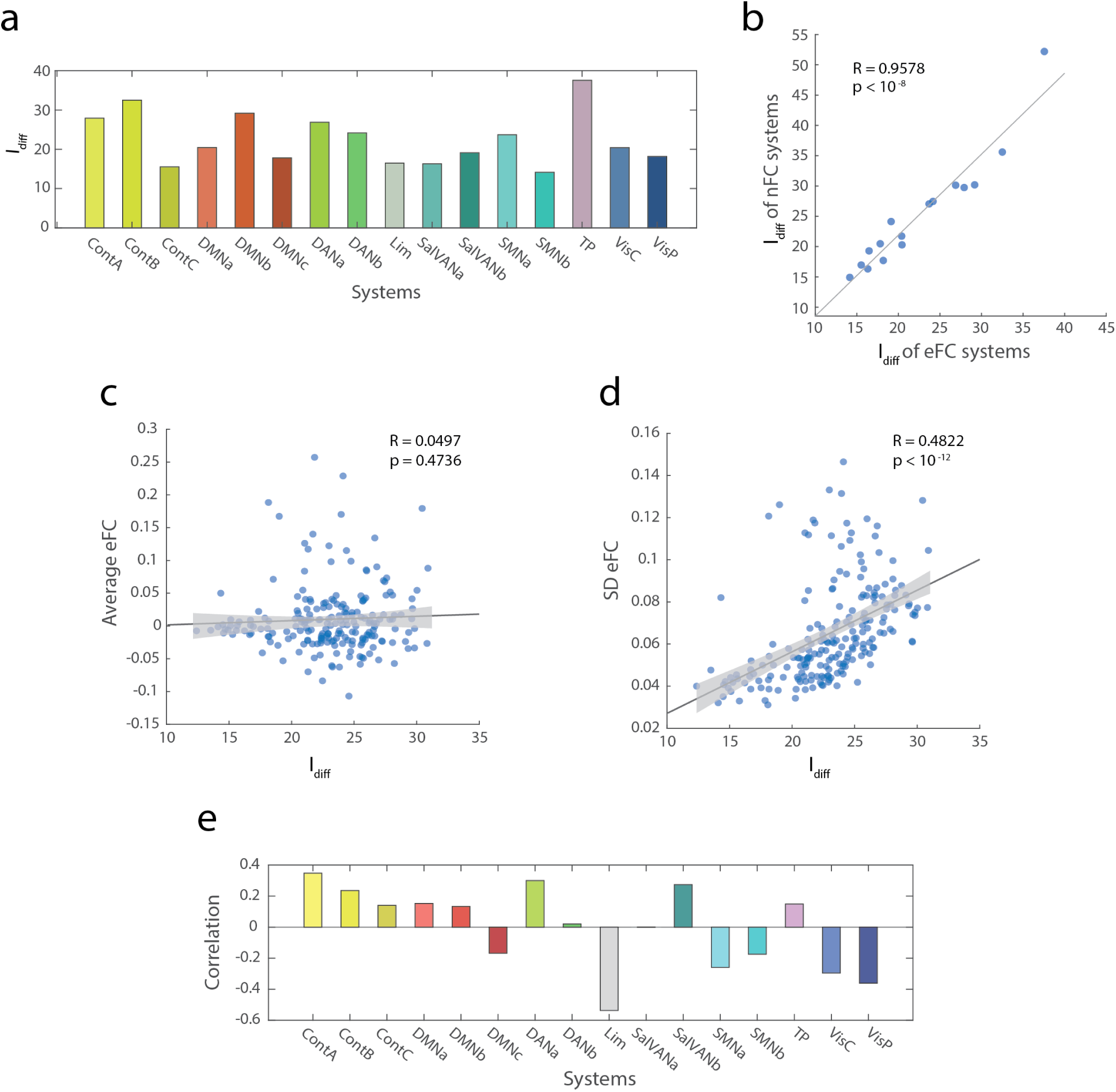
System- and cluster-level characteristics of edge functional connectivity (eFC). The within-versus between-subject similarity of eFC in the MSC dataset shows variance of identifiability across systems and high correlation to the identifiability of systems using FC. Panel (*a*) shows the within subject minus between subject similarity of eFC for systems for 10 rsfMRI scans from 10 subjects. Panel (*b*) shows the high correlation between identifiability in eFC and nFC by systems. (*c*) *I*_*diff*_ was not correlated with the average values of eFC in each cluster but in (*d*) the standard deviation of eFC values of eFC in each cluster significantly correlated with *I*_*diff*_. The bootstrapped 95% confidence interval is shaded in grey. (*e*) The correlation of 16 canonical brain systems to *I*_*diff*_ of eFC clusters.

To investigate *I*_*diff*_ of edge pairs whose nodes originate in different systems, we clustered the eFC matrices using a standard k-means algorithm, varying the number of clusters from *k* = 2 to *k* = 20. We then repeated the clustering algorithm to create a representative cluster of the dataset for each *k* and calculated the *I*_*diff*_ for each cluster. Here, we found that the mean eFC of a cluster was not correlated with its *I*_*diff*_ (*R* = 0.050, *p* = 0.474; Fig. 4*c*). On the other hand, the variability of eFC weights within a block was positively correlated with *I*_*diff*_ (*R* = 0.482, *p <* 10^−12^; Fig. 4*d*). What systems might be responsible for driving high levels of *I*_*diff*_ ? To address this question, we calculated how frequently each system was represented within a given cluster, and, separately for each brain system, calculated the correlation of this frequency with *I*_*diff*_. In general, we found that the presence of control, dorsal attention, and temporoparietal nodes in a cluster is positively correlated with it’s *I*_*diff*_. In contrast, the presence of nodes in limbic and sensorimotor systems (somatomotor and visual) are associated with reduced *I*_*diff*_ (Fig. 4*e*).

Collectively, these results suggest that in eFC, higher order cognitive systems, e.g. control, attention, and default mode networks, contribute to enhance subject identifiability, while sensorimotor and limbic networks reduce identifiability. As in the previous section, these results are in line with previous analyses using nFC demonstrating that similar systems and regions promote enhanced identifiability [14, 44]. Furthermore, our results suggest that the intrinsic heterogeneity and variability of connection weights may be an underlying factor explaining why certain systems are associated with higher or lower levels of identifiability.

### Reconstructed eFC using PCA improves subject identifiability

Throughout this manuscript, we have focused on calculating *I*_*diff*_ using the full eFC matrix or specific subsets of its edge-edge connections. As a final analysis, we wanted to test whether we could improve differential identifiability by optimally reconstructing eFC using a relatively small number of its principal components.

Previous research has used principal component analysis (PCA) of nFC to enhance identifiability [30, 35]. Briefly, this procedure entails concatenating nFC (or in this case, eFC) from all subjects and scans into a single matrix, decomposing this matrix into its principal components (PCs), and reconstructing eFC by gradually including more and more of its PCs (in ascending order of their eigenvalues). With each reconstruction, we calculated the *I*_*diff*_ with added PCs using each scan’s reconstructed eFC. Here, we apply this technique to both nFC and eFC from the MSC dataset.

Using this reconstruction method, we found that *I*_*diff*_ could be improved for both nFC and eFC. In both nFC and eFC, *I*_*diff*_ peaked at *k* = 10 components (corresponding to the number of subjects) and eFC outperformed nFC in subject identification (peak value of *I*_*diff*_ = 35.27 compared to *I*_*diff*_ = 21.17; Fig. 5*b*,*e*). Why, then, do the number of PCs match the number of subjects when optimizing for identifiability? First, we tested the effect of scans per subject on the number of PCs for maximizing identifiability. When testing for two scans per subject with 100 subjects Fig. S4*c* and with randomly selected two scans per subject with 10 subjects Fig. S4*a*, the number of PCs required for optimizing identifiability matched the number of subjects in the dataset.

**FIG. 5.**
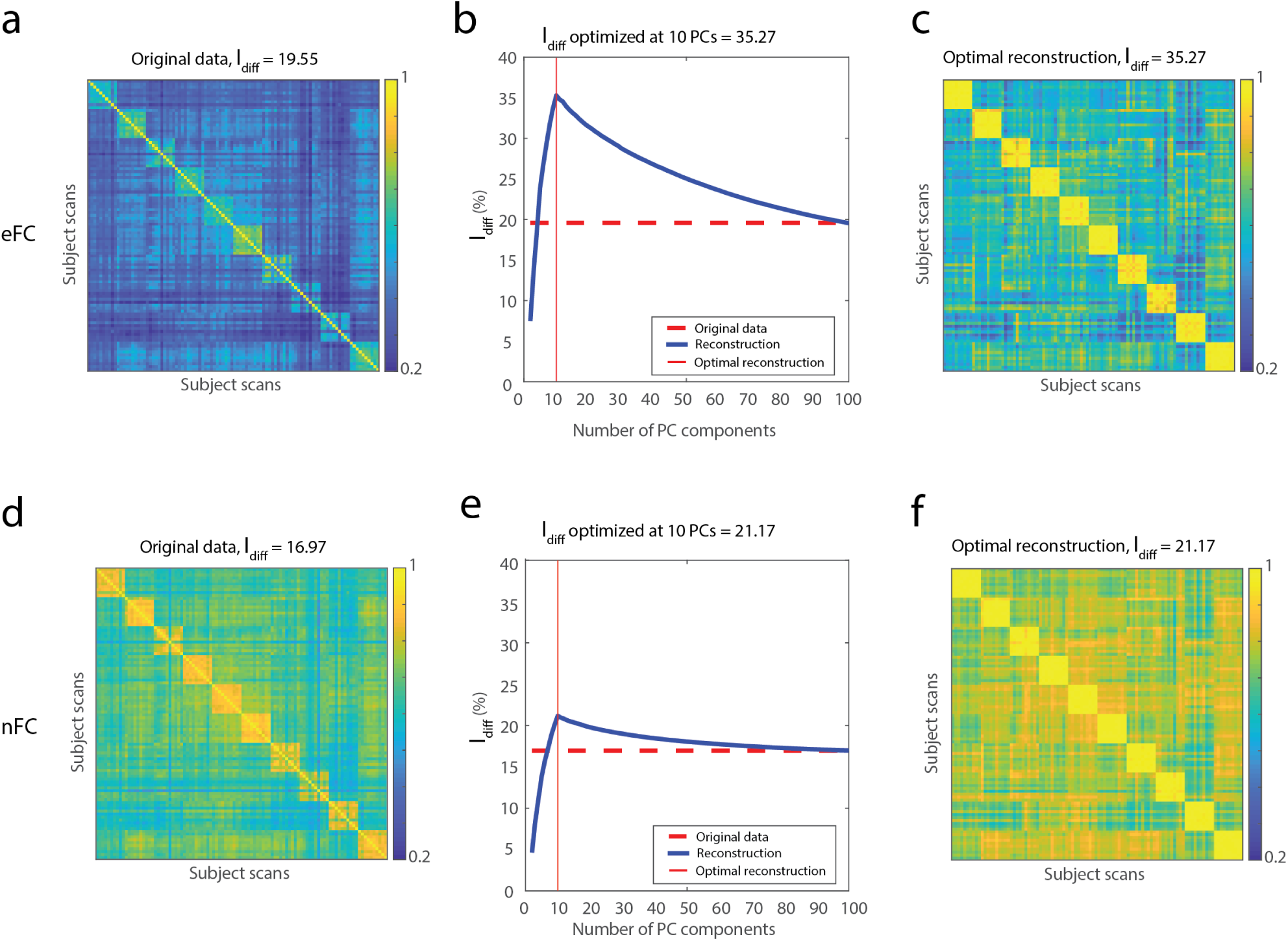
PCA reconstruction for differential identifiability optimization with eFC and FC. (*a*) Scan-to-scan similarity (Pearson correlation) matrix of eFC used as the input data for PCA. (*b*) Optimal *I*_*diff*_ measured when PC = 10; red dotted line = *I*_*diff*_ orig. (*c*) Reconstructed output eFC matrix for maximal *I*_*diff*_ using 10 PCs. (*d*) Scan-to-scan similarity (Pearson correlation) matrix of nFC used as the input data for PCA. (*e*) Optimal *I*_*diff*_ measured when PC = 10, red dotted line = *I*_*diff*_ orig. (*f*) Reconstructed output nFC matrix for maximal *I*_*diff*_ using 10 PCs.

Next, we investigated the PC coefficients that improve (PC = 1 – 10) or reduce (PC = 11 – 100) identifiability. The first prinicipal component (PC1), mathematically, explains the largest variance of eFC values across scans and subjects. PC1 was the only component whose coefficients were uniformly positive (Fig. S5). The next nine coefficients (PC2 – PC10) expressed “blocky” patterns that correspond to single subjects (Fig. S5), while this pattern was absent in PC11 – PC100 (Fig. S6).

In agreement with previous reports [35, 47], our results demonstrate that subject identification can be improved by selectively retaining a subset of components that match the number of subjects in the dataset. We show that the magnitude of improvement is considerably greater using eFC compared to nFC, suggesting that eFC may better capture personalized and idiosyncratic features compared to nFC [31, 48].

## DISCUSSION

Here, we applied subject identification to a novel, edge-centric network representation of the human cerebral cortex. We found that given sufficient scan length, eFC exhibits greater levels of differential identifiability than nFC, an improvement that we linked to contributions made by brain regions in association cortex. Finally, we used a dimension-reduction and reconstruction method to show that the relative improvement in identifiability enjoyed by eFC could be further enhanced, highlighting the potential for eFC to be used in future studies.

### Edge functional connectivity enhances subject identifiability

Central to this paper is the observation that eFC results in improved subject identification relative to conventional node-based connectivity, nFC. Whereas nFC measures the similarity of activity between two brain regions – a first-order correlation [49] – eFC measures the similarity of co-fluctuations between edge pairs – a higher-order correlation [36]. Understanding the higher-order organization of networks has proven useful in other disciplines [40, 41, 50, 51]. Here, adopting an edge-centric perspective allows us to link higher-order brain network organization with subject specific features.

We asked whether the eFC – a higher-order reconstruction – leads to improved identification of subjects. That is, if we were to examine the identifiability of whole brain nFC and eFC, would eFC allow us to more accurately identify a single subject based on their connectivity data? We also asked whether identification using eFC would be impacted by scan length and amount of data? In general, we found that given sufficient amounts of data, eFC outperformed nFC in terms of identifiability. These results held across two datasets and two different parcellations, suggesting that higher-order network structure carries important subject-specific information and that idiosyncratic features of networks are hence better captured *via* an edge-centric viewpoint. These observations both challenge and expand current knowledge of subject identification and precision network mapping [14, 15, 28, 35, 48]. Future investigation is required for understanding the relationship between scan length and higher-order brain representations [49].

### Heteromodal regions drive subject identification

Which brain regions drive the subject specificity of eFC? Are there particular regions of the brain that make a subject more or less identifiable? To answer these questions, we analyzed subject identifiability on three different scales. First, we analyzed each node’s contribution to *I*_*diff*_ using a “leave-one-node-out” approach. We found that when removing certain nodes prior to constructing the eFC matrices, resulted in significantly reduced or increased *I*_*diff*_. In particular, nodes and edges from higher-order systems lead to significant reductions in *I*_*diff*_, while those associated with sensorimotor and limbic regions yielded increased identifiability when removed prior to eFC construction. These findings support earlier findings in nFC in which brain regions supporting higher-order function drive subject identification compared to regions of unimodal function [14]. We suspect the overlapping character of heteromodal versus unimodal brain regions as an explanation of this result. Future research can answer questions such as whether inter-subject idiosyncrasies and brain regions involved are modulated by the fMRI’s task.

Next, we asked whether higher and lower *I*_*diff*_ is attributable to edge pairs from a single canonical brain system [46]. To address this question, we estimated subjects’ eFC matrices separately from edge pairs constructed using only one of sixteen brain systems. We found that identifiability from single-system eFC and nFC were significantly positively correlated and that edges forming stubs from heteromodal brain regions tended to have higher *I*_*diff*_. In other words, the cohesive edge pairs within single systems are likely not driving eFC’s improved *I*_*diff*_ observed globally. Rather, it suggests that edges falling between different brain networks may be driving the improved *I*_*diff*_ in eFC compared to that of nFC.

The number of vertices and connections in eFC and nFC matrices diverge by orders of magnitude. To match their dimensionality and ensure a fairer comparison, we clustered nFC and eFC matrices into the same number of communities. We found that the variance of connections within eFC clusters was positively correlated with subject identification, which the average weight of those connections were not. We also found that clusters containing edges whose stubs originated in heteromodal systems resulted in greater levels of identifiability compared to clusters composed of edges associated with unimodal brain systems. These results are in agreement with findings from nFC studies [14, 45] and with more recent studies involving clusters of eFC [38]. Our findings suggest that variability of connection weights within systems and clusters may be an important feature driving the efficacy of connectome fingerprinting and identifiability.

Collectively, our findings suggest that subject identification is driven by heteromodal regions from higher-order brain systems. These observations have clear implications for generating robust network biomarkers with subject-specific information, while reducing the amount of required data to its subset. More importantly, our results demonstrate that clusters of eFC with high variance, which maybe undermined in group-level analyses, may be useful in determining subject-specific characteristics or in personalized medical treatment. Also, this study leaves clues for future research on task-specific biomarkers to further specify the effect of tasks during fMRI acquisition on identifiability and to maximize subject-specific characteristics without requiring scans of prohibitive length, especially for vulnerable and clinical populations [12]. Lastly, clustering eFC matrices showed potential as a method of dimensionality reduction that is robust across subjects. Future work is necessary to clarify clusters or subsets of edges that are robust for group-level and subject-level brain networks.

### Principal Component Analysis highlights idiosyncrasies in eFC

Here, we followed recent nFC studies and applied PCA to eFC data, effectively reducing eFC to a small set of principal components [30, 35]. We found that selectively reconstructing eFC using only those components that explained the greatest variance resulted in improved subject identifiability, far beyond the improvements when an identical procedure was applied to nFC data. This suggests that given the exactly same fMRI BOLD data, we can extract enhanced subject-level fingerprints from eFC data.

Interestingly, the number of PCs required to optimize eFC’s subject identifiability matched the number of subjects in two independently acquired datasets. These results parallel previous research using nFC for subject identification [35]. Our results additionally show that this is robust despite a single subject having more than one test-retest pair of rsfMRI scans. Why, then, do the number of PCs match the number of subjects when optimizing for identifiability? To address this question, we analyzed the coefficients of PCs that increase (PC = 1-10) versus those that decrease (PC = 11-100) identifiability. The first principal component, mathematically, should explain the largest variance of eFC values across scans. This was the only PC out of 100 that had a consistently positive value. Also, only PCs 2 to 10 showed significant “blockiness” for specific subject’s scans. One possibility is that the first PC explains group-level eFC variance, the underlying group-level eFC features, whereas PCs 2 to 10 tend to explain subject-level eFC variance. We speculate that the number of blocky PCs are *N −* 1 (N = number of subjects) since every subject can be identified with *N −* 1 PCs *via* the process of elimination. These results provide further evidence that eFC can be a valuable framework when investigating and improving subject identifiability with linear transformation algorithms such as PCA. However, future work is necessary to disseminate the precise features of these PCs and alternative dimensionality reduction methods such as factor analysis or CCA [52, 53] should be explored for *I*_*diff*_ optimization.

### Future directions

Our results present exciting possibilities for future studies. Here, we used a previously-defined measure of identifiability. However, this measure can be misleading in some cases. For instance, *I*_*diff*_ can still take on high values if most subjects exhibit high levels of self-similarity, even if the remaining subjects exhibit poor self-similarity [27, 35, 54, 55]. Future work should investigate alternative measures for quantifying the performance of subject identification procedures.

Other possibilities for future studies include applying predictive modeling to eFC and delineating those components of eFC that maximize subject idiosyncrasies while reducing the amount of data required [56]. In addition, similar optimization approaches could be used to study individual differences in cognitive, developmental, and disease states based on features extracted from eFC [22].

### Limitations

An overarching limitation associated with eFC is that linking it back to individual brain regions is challenging. Each connection in the eFC matrix always involves four nodes (two edge-edge connections), and allocating its properties to any one brain region or cognitive system is, in most cases, not possible. Here, we circumvented this complication by measuring the effect of a node’s removal (equivalent to 199 edges in a 200 node parcellation) from the resulting eFC’s identifiability and only accounting for edge pairs derived from single systems. While these approaches attempt to locate edges to brain regions, future work is necessary to determine a robust method in tracing eFC elements back to specific locations in brain.

A second limitation concerns the measure of “differential identifiability,” which accounts for both within-subject and between-subject similarity. However, we acknowledge alternative measures of identifiability such as calculating the subject identification accuracy of FC-FC correlations [14], ROC (receiver operating characteristic) curve accuracies [27], and using a Nearest Centroid Classification for each subject in graph embedding representations[55]. Nonetheless, we focused our analyses using *I*_*diff*_ since this approach methodologically extends a popular identification approach [14] while accounting for “common features” shared across subjects [35]. However, other methods for measuring identifiability and comparing fMRI data should also be investigated and developed to further understand idiosyncrasies in edge-centric FC.

A final limitation concerns the procedure for estimating edge clusters and utilizing its results. Here, we use a k-means algorithm to partition edges into a group-wise fixed number of clusters based on the eFC similarity. The benefit of this approach to estimating edge clusters is that this algorithm is computationally efficient and can be calculated using a distance metric. Given the extensive list of alternative methods for clustering [57–59], other algorithms must be investigated for detecting edge clusters. Also, between-cluster edges may act as bridges between functional systems that further explain idiosyncrasies and behavioral associations [38]. However, we add that our analyses exclude between-cluster edges and focus on within-cluster edges. Future studies should investigate the effects of varying clustering algorithms and study the effect of within-cluster versus between-cluster edges on individual differences.

## MATERIALS AND METHODS

### Datasets

The Midnight Scan Club (MSC) dataset [15] included rsfMRI from 10 adults (50 female, age = 29.1 *±* 3.3). The study was approved by the Washington University School of Medicine Human Studies Committee and Institutional Review Board with the informed consent from all subjects. 12 scans from each subject were acquired on separate days starting from midnight. Per each subject, 10 rsfMRI scans were collected with a gradient-echo EPI sequence(run duration = 30 min, TR = 2200ms, TE = 27ms, flip angle = 90°) with the participant’s eyes open while recording eyetracking to monitor for prolonged eye closure (for assessing drowsiness). Images were collected on a 3T Siemens Trio.

The Human Connectome Project (HCP) dataset [60] included resting state functional data (rsfMRI) from 100 unrelated adult subjects (54% female, mean age = 29.11 *±* 3.67, age range = 22-36).These subjects were selected as they comprised the “100 Unrelated Subjects” (U100) released by the Human Connectome Project. The study was approved by the Washington University Institutional Review Board and informed consent was obtained from all subjects. Subjects underwent four 15 minute rsfMRI scans over a two day span. A full description of the imaging parameters and image preprocessing can be found in [61]. The rsfMRI data was acquired with a gradient-echo EPI sequence (run duration = 14:33 min, TR = 720ms, TE = 33.1ms, flip angle = 52°, 2mm isotropic voxel resolution, multiband factor = 8) with eyes open and instructions to fixate on a cross. Images were collected on a 3T Siemens Connectome Skyra with a 32-channel head coil.

### Image Preprocessing

#### MSC Functional Preprocessing

Functional images in the MSC dataset were pre-processed using *fMRIPrep* 1.3.2 [62], which is based on Nipype 1.1.9 [63]. The following description of *fMRIPrep*’s preprocessing is based on boilerplate distributed with the software covered by a “no rights reserved” (CCO) license. Internal operations of *fMRIPrep* use Nilearn 0.5.0 [64], ANTs 2.2.0, FreeSurfer 6.0.1, FSL 5.0.9, and AFNI v16.2.07. For more details about the pipeline, see the section corresponding to workflows in *fMRIPrep*’s documentation.

The T1-weighted (T1w) image was corrected for intensity non-uniformity with N4BiasFieldCorrection [65, 66], distributed with ANTS, and used as T1w-reference throughout the workflow. The T1w-reference was then skull-stripped with a Nipype implementation of the ants-BrainExtraction.sh workflow, using NKI as the target template. Brain surfaces were reconstructed using recon-all [67], and the brain mask estimated previously was refined with a custom variation of the method to reconcile ANTs-derived and FreeSurfer-derived segmentations of the cortical gray-matter using Mindboggle [68]. Spatial normalization to the *ICBM 152 Nonlinear Asymmetrical template version 2009c* [69] was performed through nonlinear registration using antsRegistration, using brain-extracted versions of both T1w volume and template. Brain tissue segmentation of cerebrospinal fluid (CSF), white-matter (WM) and graymatter(GM) was performed on the brain-extracted T1w using FSL’s fast [70].

Functional data was slice time corrected using AFNI’s `3dTshift` and motion corrected using `FSL’s mcflirt`. *Fieldmap-less* distortion correction was performed by co-registering the functional image to the same subject. T1w image with intensity inverted constrained with an average fieldmap template, implemented with `antsRegistration`. This was followed by co-registration to the corresponding T1w using boundary-based registration with 9 degrees of freedom. Motion correcting transformations, field distortion correcting warp, BOLD-to-T1w transformation and T1w-to-template (MNI) warp were concatenated and applied in a single step using `antsApplyTransforms` using Lanczos interpolation. Several confounding timeseries were calculated based on this preprocessed BOLD framewise displacement (FD), DVARS and three region-wise global signals. FD and DVARS are calculated for each functional run, both using their implementations in Nipype. The three global signals are extracted within the CSF, the WM, and the whole-brain masks. The resultant NIFTI file for each MSC subject used in this study followed the file naming pattern: `* spaceT1w descpreproc bold.nii.gz`.

#### HCP Functional Preprocessing

Functional images in the HCP dataset were minimally preprocessed according to the description provided in [61]. Briefly, these data were corrected for gradient distortion, and motion, and then aligned to a corresponding T1-weighted (T1w) image with one spline interpolation step. This volume was further corrected for intensity bias and normalized to a mean of 10000. This volume was then projected to the 32k fs LR mesh, excluding outliers, and aligned to a common space using a multi-modal surface registration [71]. The resultant CIFTI file for each HCP subject used in this study followed the file naming pattern: `* REST{1,2} {LR,RL} Atlas MSMAll.dtseries.nii`.

#### Image Quality Control

All functional images in the MSC and HCP datasets were retained. The quality of functional images in the MSC were assessed using fMRIPrep’s visual reports and MRIQC 0.15.1 [62]. MSC data was visually inspected for whole brain field of view coverage, signal artifacts, and proper alignment to the corresponding anatomical image. All time series data were visually inspected as well.

### Functional Networks Preprocessing

#### Parcellation Preprocessing

A functional parcellation designed to optimize both local gradient and global similarity measures of the fMRI signal[46] (Schaefer200) was used to define 200 areas on the cerebral cortex. These nodes are also mapped to the Yeo canonical functional networks [72]. For the HCP dataset, the Schaefer200 is openly available in 32k fs LR space as a CIFTI file. For the MSC dataset, a Schaefer200 parcellation was obtained for each subject using a Gaussian classifier surface atlas[73] (trained on 100 unrelated HCP subjects) and FreeSurfer’s mris ca abel function. These tools utilize the surface registrations computed in the recon-all pipeline to transfer a group average atlas to subject space based on individual surface curvature and sulcal patterns. This method rendered a T1w space volume for each subject. For use with functional data, the parcellation was resampled to 2mm T1w space. This process could be repeated for other resolutions of the parcellation (i.e. Schaefer100).

#### Functional Network Preprocessing

Each preprocessed BOLD image was linearly detrended, band-pass filtered (0.008 – 0.08 Hz) [74], confound regressed and standardized using Nilearn’s signal.clean, which removes confounds orthogonally to the temporal filters [75]. The confound regression employed [76] included 6 motion estimates, time series of the mean CSF, mean WF, and mean global signal, the derivatives of these nine regressors, and the squares of these 18 terms. Furthermore, a spike regressor was added for each fMRI frame exceeding a motion threshold (MSC = 0.5mm framewise displacement, HCP = 0.25mm root mean squared displacement). This confound strategy has been shown to be a relatively effective option for reducing motion-related artifacts [74]. Following preprocessing and nuisance regression, residual mean BOLD time series at each node were recovered.

### Edge graph construction

eFC can be calculated by acquiring the regional time series data and their z-scores. Next, for all pairs of brain regions, we calculate the element-wise product of their z-scored time series. This returns the “edge time series” that represent the magnitude of co-fluctuation for pairs of brain regions which can be correlated across time as the Pearson correlation coefficient. Lastly, the scalar product between pairs of edge time seires is calculated and repeated over all pairs of edges to create an edge-by-edge matrix, which are normalized to the interval [-1, 1].

### Differential Identifiability

The functional connectome’s identifiability or finger-printing is based on the assumption that a single subject’s connectivity profile should be, more similar within the same subject across scans and sessions than between different subjects. Previous research using the conventional functional connectome [14] showed that, robust identification of an individual is possible using sample FC to find the “target” FC of the subject in a pool of subject FCs with Pearson correlation analyses. Prior research on quantifying individual differences in functional connectivity include calculating the *geodesic distance* [77] *and Pearson correlation* across individual’s scans [35]. While the geodesic distance approach also provides a summary measure of the inter-scan differences, we adopt the quantification metric by Amico et al. [35], which takes into consideration the covariance and standard deviation of the eFC and FC matrices. This metric is called the differential identifiability (*I*_*diff*_) derived from the “identifiability matrix”, i.e. the matrix of correlations (Pearson) between subjects’ FCs. The *I*_*diff*_ is calculated by quantifying self-identifiability or *I*_*self*_ and substracting between subject similarity or *I*_*others*_, represented as the diagonal and off-diagonal elements of the identifiability matrix (Fig.1b). Differential identifiability (*I*_*diff*_) of a group of subjects can be summarized as the following:

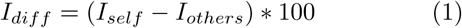

which is the difference of average within-subject similarity and average between-subjects similarity of FCs. A high value of *I*_*diff*_ in a dataset will have higher subject identifiability and the optimization of identifiability is reduced to maximizing *I*_*diff*_.

To test for reproducibility and the effect of parcellations, we repeated the analysis with 100 unrelated subjects from the HCP dataset [60]. Both datasets were tested for effects of numbers of parcellations by measuring *differential identifiability* in 100 and 200 node datasets (Fig. S1).

### Tracing identifiability to brain regions by a leave-one-node-out approach

In the previous section, we described the procedure for calculating *I*_*diff*_, a measure known to quantify the relative similarity of scans within versus between subjects[43]. In this section, we test which brain regions reduce subject identification when removed from the calculation of eFC, compared to when measuring *I*_*diff*_ with the entire eFC matrix. In addition, we test whether removal of specific brain systems significantly reduce *I*_*diff*_ compared to using the whole-brain eFC.

The direct connection between eFC’s edge-edge pair co-fluctuation and a brain region can somewhat be an arbitrary procedure since there can be up to 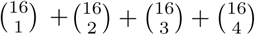 combinations of brain regions in a single edge pair in eFC. To avoid making assumptions that edge pairs’ weights are linearly related to the edge pair strength, we adopt a leave-one-node-out approach prior to eFC matrix construction. The effect of removing a node prior to eFC construction was calculated by subtracting the *I*_*diff*_ of the single-node-removed eFC from the whole-brain *I*_*diff*_ (Fig. S3*a-b*).

Next, we examined the effect of each functional brain system, and if removed, the difference in *I*_*diff*_ measured from eFC. Analogous to the single node removal approach earlier, we removed single systems (i.e. all nodes from a single system) and measured the effect on *I*_*diff*_ by subtracting this result from the whole-brain *I*_*diff*_ (Fig. 3*c-d*). We did not control for the difference in number of nodes (i.e. entire brain = 200 nodes, single-system-removed *≈* 190 nodes) since the effect of the total number of nodes to *I*_*diff*_ is unclear.

### Single system edge pairs identifiability

The benefit and caveat of the leave-one-node-out approach is that it removes all edge pairs involving a particular node or system due to eFC’s overlapping characteristic[36]. Therefore, we were still unclear of the effects of a purely single system due to this characterisic. In order to determine the *I*_*diff*_ a single-system, we extracted the edge pairs that include nodes only from a particular system4*a* and measured the *I*_*diff*_. The analogous approach was applied to nFC, which included node-pairs from the nFC which include nodes from a single system. The single-system level node-pairs were compared to that of edge pairs from eFC (Pearson correlation). Also, subject similarity matrices using edge pairs from single systems are included in (Fig. S2).

### K-means clustering for identifiability

The eFC matrices used here have approximately a squared dimensionality of components compared to the conventional nodal FC matrices. While the higher dimensionality of eFC may provide insight into the relationship of it’s components (the edge-edge communications) that is not directly shown in FC matrices, clustering the components of the eFC matrices present a computational challenge. This is especially the case if the number of partitions of the matrix is unknown and is left for exploration. To address this issue and cluster the eFC, we have applied a simple two-step clustering procedure that operates on a low-dimensional representation of the eFC matrix.

First, we performed an eigendecomposition of the eFC matrix, retaining the top 50 eigenvectors. These eigen-vectors were rescaled to the interval [-1, 1] by dividing each eigenvector by its largest magnitude element. Then we simply clustered the rescaled eigenvectors using a standard k-means algorithm with Euclidean distance. We varied the number of communities, *k*, from *k* = 2 to *k* = 20, repeating the clustering algorithm 250 times at each values. We retained as a representative partition, the one with the greatest overall similarity to all other partitions.

In our third analysis, we observed clusters of *k* = 2 to 20 using a standard k-means clustering. We found clusters’ *I*_*diff*_ to positively correlate with variance in eFC but found no correlation with the average eFC. Also, our results reveal clusters with higher *I*_*diff*_ were composed of nodes in association brain regions rather than nodes of sensorimotor, limbic brain regions. Combined, our results suggest that by capitalizing on clusters which convey a more diverse repertoire of eFC values and with nodes from association systems, we may generate clusters, or subsets of edge pairs, that robustly maximize subject identification while reducing the amount of required data[29].

### Principal Component Analysis

Principal component analysis (PCA) is a widely used statistical method [78] that allows exploration of the underlying structure of the data. PCA transforms a set of observed data with potentially correlated variables into a set of linearly uncorrelated variables called principal components. These principal components are then ranked in a descending order that explains the most to least variance of the data. We adopted principal component analysis to directly compare eFC’s identifiability performance to that of nFC, which has been explored by Amico et al. [35].

First, the number of principal components are matched with the number of functional connectomes of the dataset. This allows for the decomposition from PCA, by definition, to account for 100% of the variance in the data. The PCs from PC = 2 to 20 were ranked by their explained variance in a descending order. Individual’s nFC and eFC were then reconstructed as a function of the number of components included based on the rationale that group-level information is carried in high variance components and subject-level information is conveyed in less higher variance components. Therefore, after extracting the main principal components, each individual’s connectivity matrix were reconstructed based on the mean and linear recombination of a select number of PCs that maximized *I*_*diff*_.

Next, we controlled for the effect of number of scans per individual, which affects the total scan duration or amount of data per subject (Fig. 2*a-b*). From the MSC dataset, we randomly selected two out of ten scans per subject as the test-retest scans for the PCA-derived *I*_*diff*_ maximization with 1000 iterations. In both eFC and nFC, *I*_*diff*_ optimized with ten PCs, which match the number of subjects in the dataset. Also, we randomly sampled 10 subjects out of 100 to test if *I*_*diff*_ optimizes at the number of subjects regardless of the individuals. From the HCP dataset, we randomly selected ten subjects out of 100 with two scans per subject. In each of the 1000 iterations, the *I*_*diff*_ for each PC was plotted for visualization (Fig. S3*c-d*).

In both the MSC and HCP dataset, we found the number of subjects to match the number of principal components in which *I*_*diff*_ is maximized in the reconstructed eFC and nFC matrices. To determine the potential driver of this result, we decomposed each PC’s coeffcients for each scan Fig. S5, Fig. S6, Fig. S7, Fig. S8. For PCs 2 to 10, each subjects’ coefficients were tested against that of the other subjects’ (*t − test* with Bonferroni-correction).

## AUTHOR CONTRIBUTIONS

YJ and RFB conceived of study, carried out all analyses, and wrote the first draft of the manuscript. JF processed the data and FZE, and OS contributed to project direction *via* discussion. All authors helped revise and write the submitted manuscript.

## DATA AVAILABILITY

All imaging data come from publicly-available, open-access repositories.Human connectome project data can be accessed *via* https://db.humanconnectome.org/app/template/Login.vm after signing a data use agreement. Midnight scan club data can be accessed *via* OpenNeuro at https://openneuro.org/datasets/dx000224/versions/1.0.1.

## CODE AVAILABILITY

All processing and analysis code is available upon reasonable request.

---

**FIG. S1.**
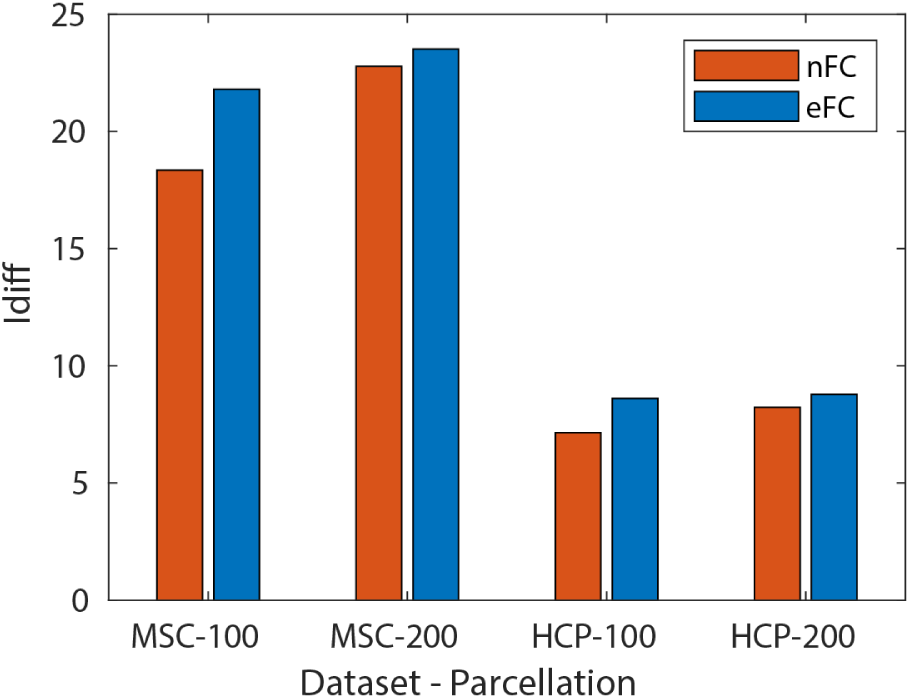
Differential identifiability (%) across varying number of nodes across datasets. *Differential identifiability* from nFC versus eFC was compared across two independently acquiured datasets – Midnight Scan Club (MSC) and Human Connectome Project (HCP) – using 100 and 200 node parcellations.

**FIG. S2.**
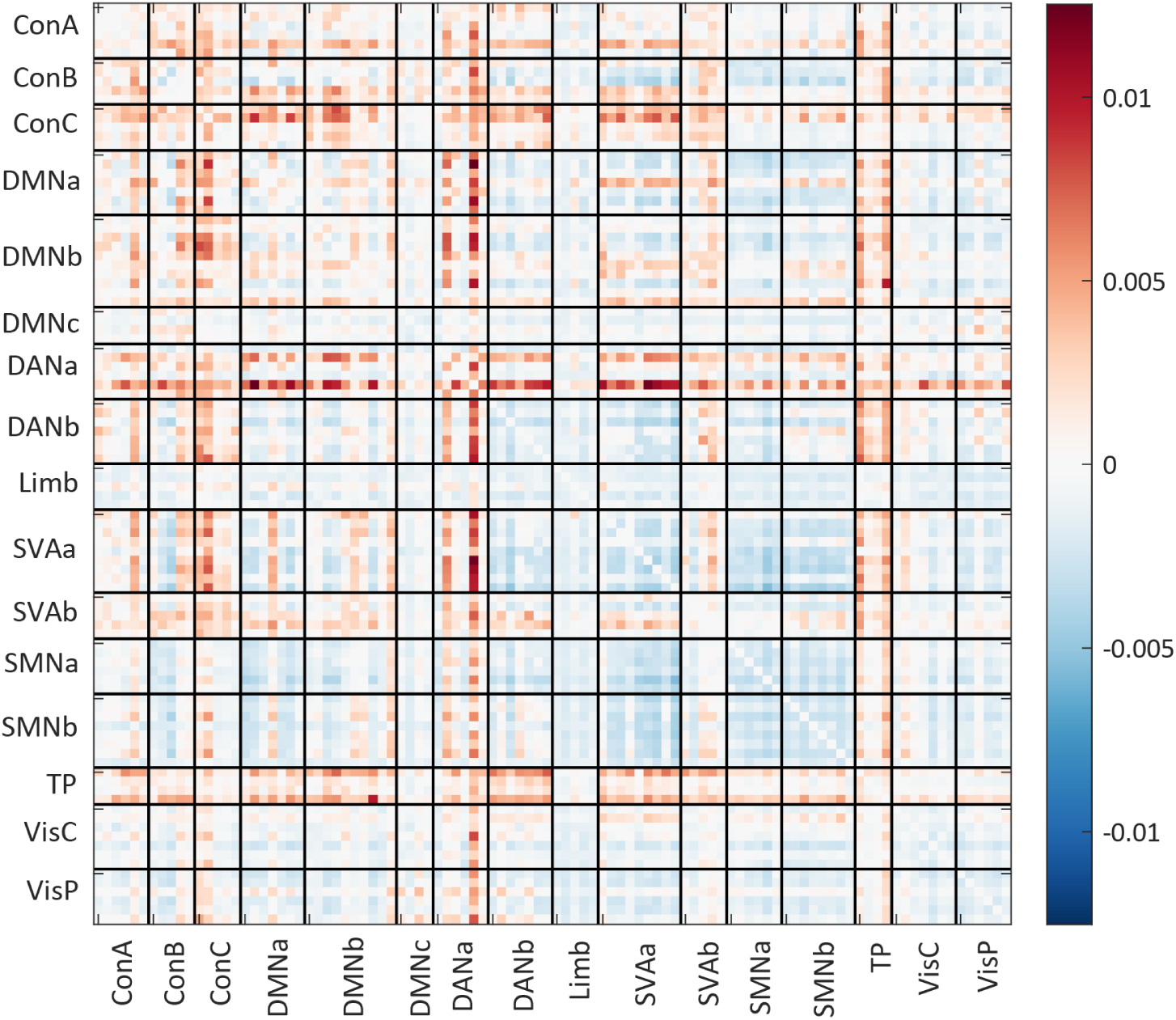
− Δ*I*_*diff*_ of “leave-one-edge-out” analysis. Relative *I*_*diff*_ post removal of an edge from each scan’s eFC matrix. Edges from the eFC matrices have been re-ordered by the corresponding stub’s system

**FIG. S3.**
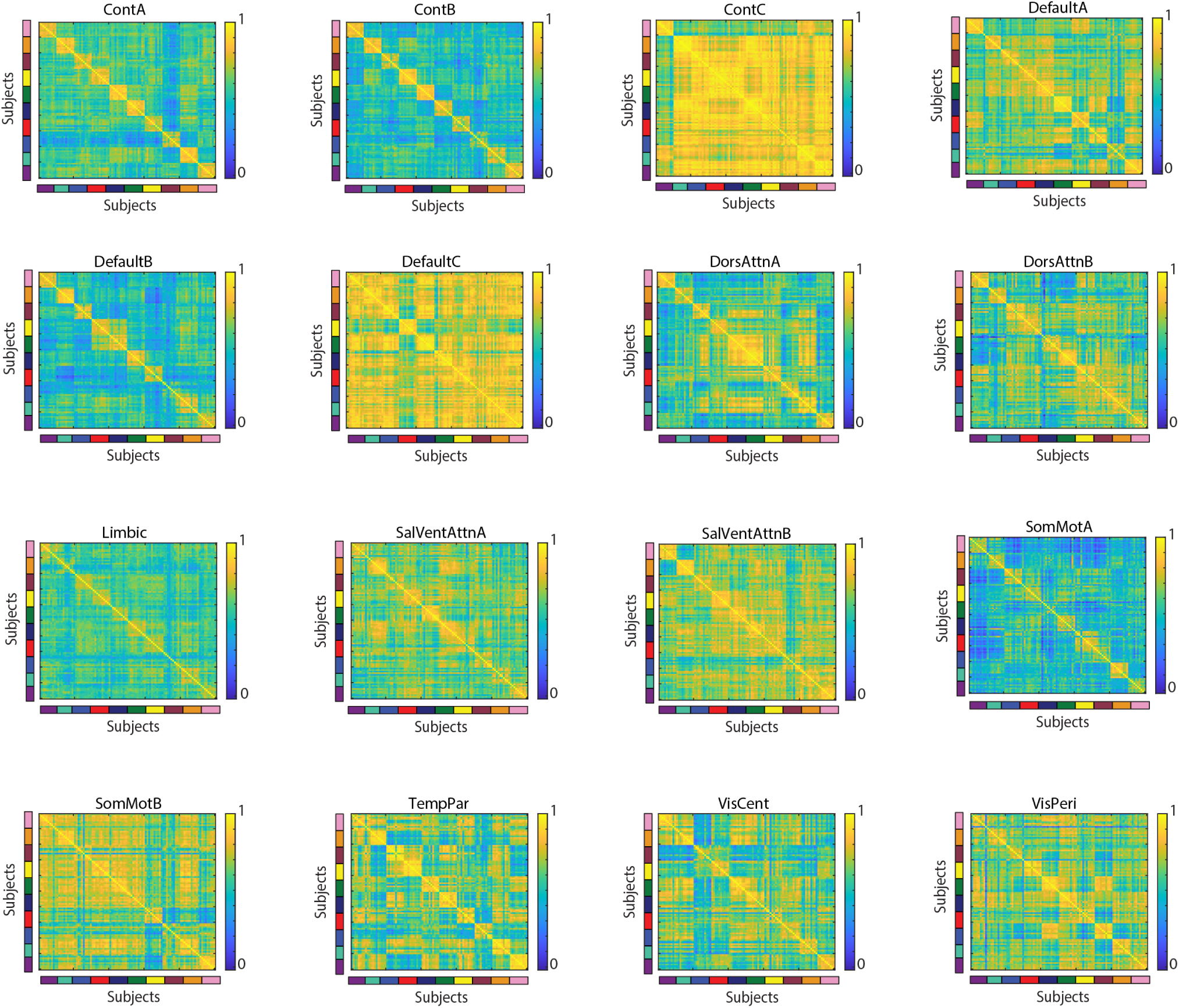
Subject similarity matrix from single-system edge pairs. Subject similarity matrices calculated by Pearson correlations of single-system derived edge pairs (all 4 nodes from a single system) from eFC.

**FIG. S4.**
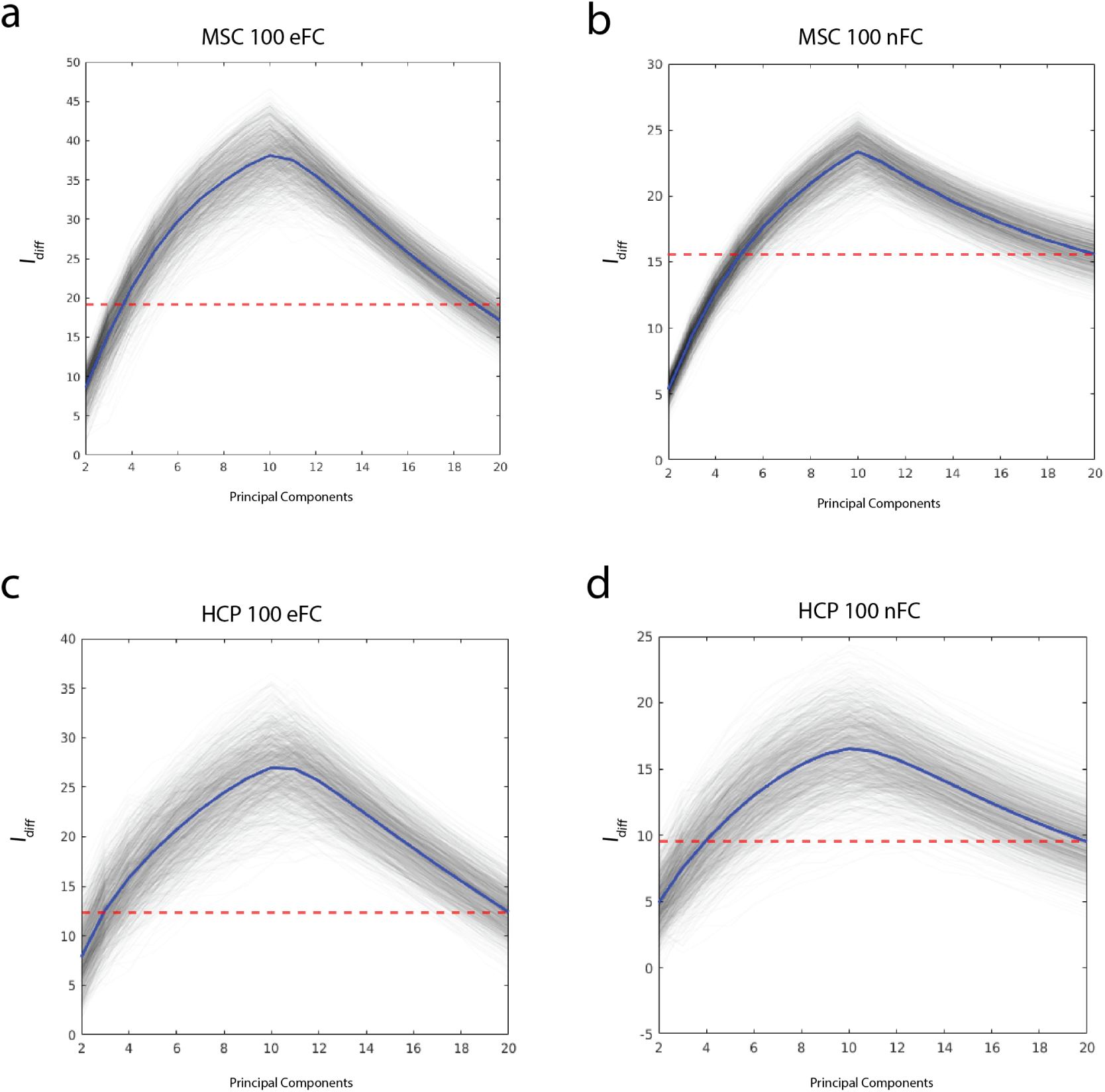
Differential identifiability (%) from reconstructed matrices with varying numbers of principal components and number of scans per subject. (*a) I*_*diff*_ across PC = 2 ∼ 20 in eFC of randomly selecting two scans per subject from the MSC dataset. (*b*) *I*_*diff*_ across PC = 2 ∼ 20 in nFC of randomly selecting two scans per subject from the MSC dataset. (*c*) *I*_*diff*_ across PC = 2∼ 20 in eFC of randomly selecting ten subjects (two scans per subject; HCP). (*d*) *I*_*diff*_ across PC = 2 ∼ 20 in nFC of randomly selecting ten subjects (two scans per subject; HCP). All panels display 100 iterations of randomization. Red-dotted lines indicate the *I*_*diff*_ from the original matrices; blue lines indicate the average *I*_*diff*_ across randomly permuted iterations; black lines indicate the *I*_*diff*_ for each PC per randomly permuted iteration.

**FIG. S5.**
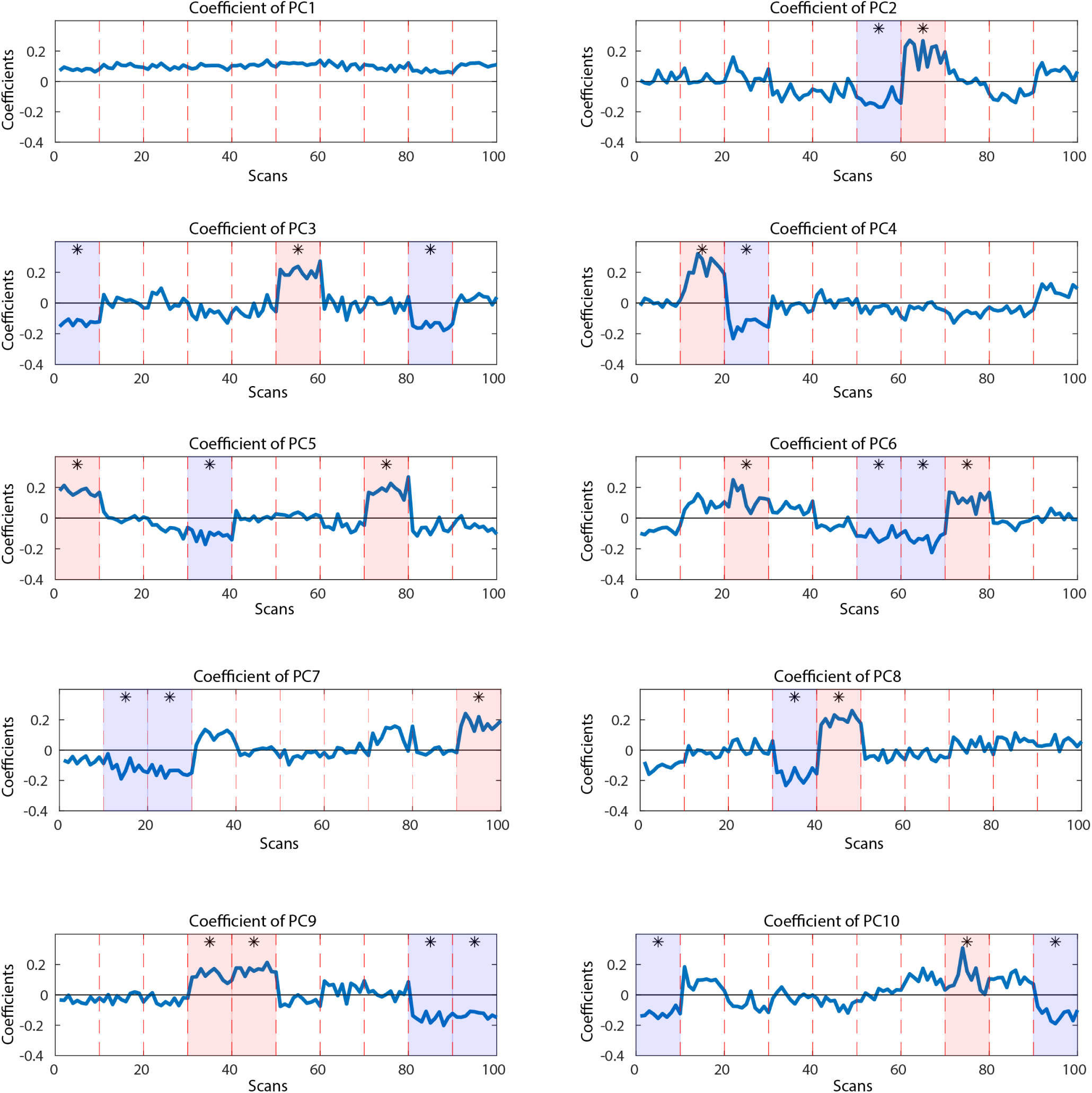
Coefficients for Principal Components 1 to 10 in MSC 100 node eFC. X-axes for all subplots are coefficients for each scan and Y-axes are the coefficients. Red-dotted lines indicate separation for each subject (10 scans per subject) and black lines indicate where coefficient equals zero. Asterisks on coefficients are for coefficients with *p <* 0.0005 (Bonferroni-corrected).

**FIG. S6.**
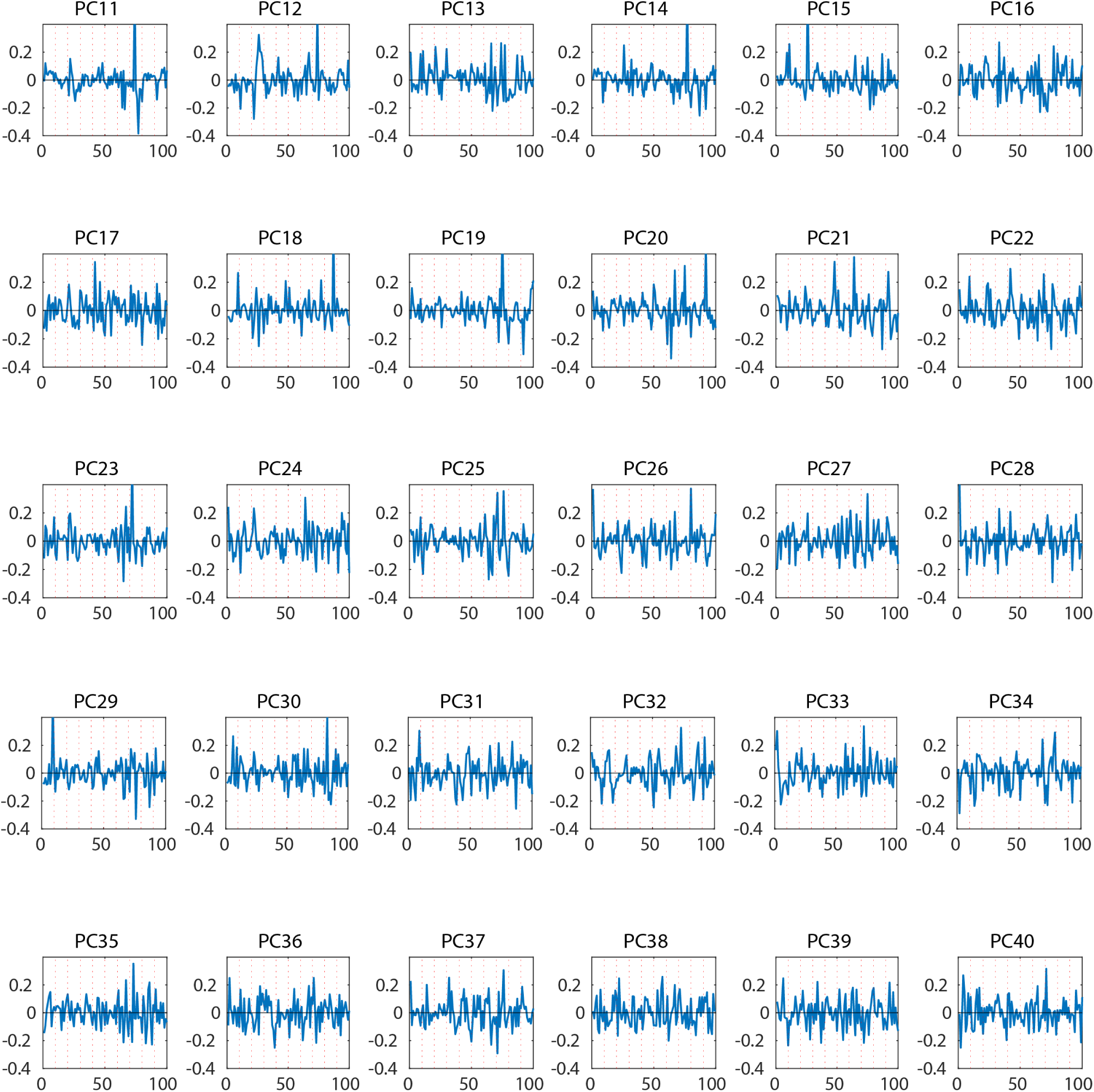
Coefficients for Principal Components 11 to 40 in MSC 100 node eFC. Red-dotted lines used for separating subjects’ scans.

**FIG. S7.**
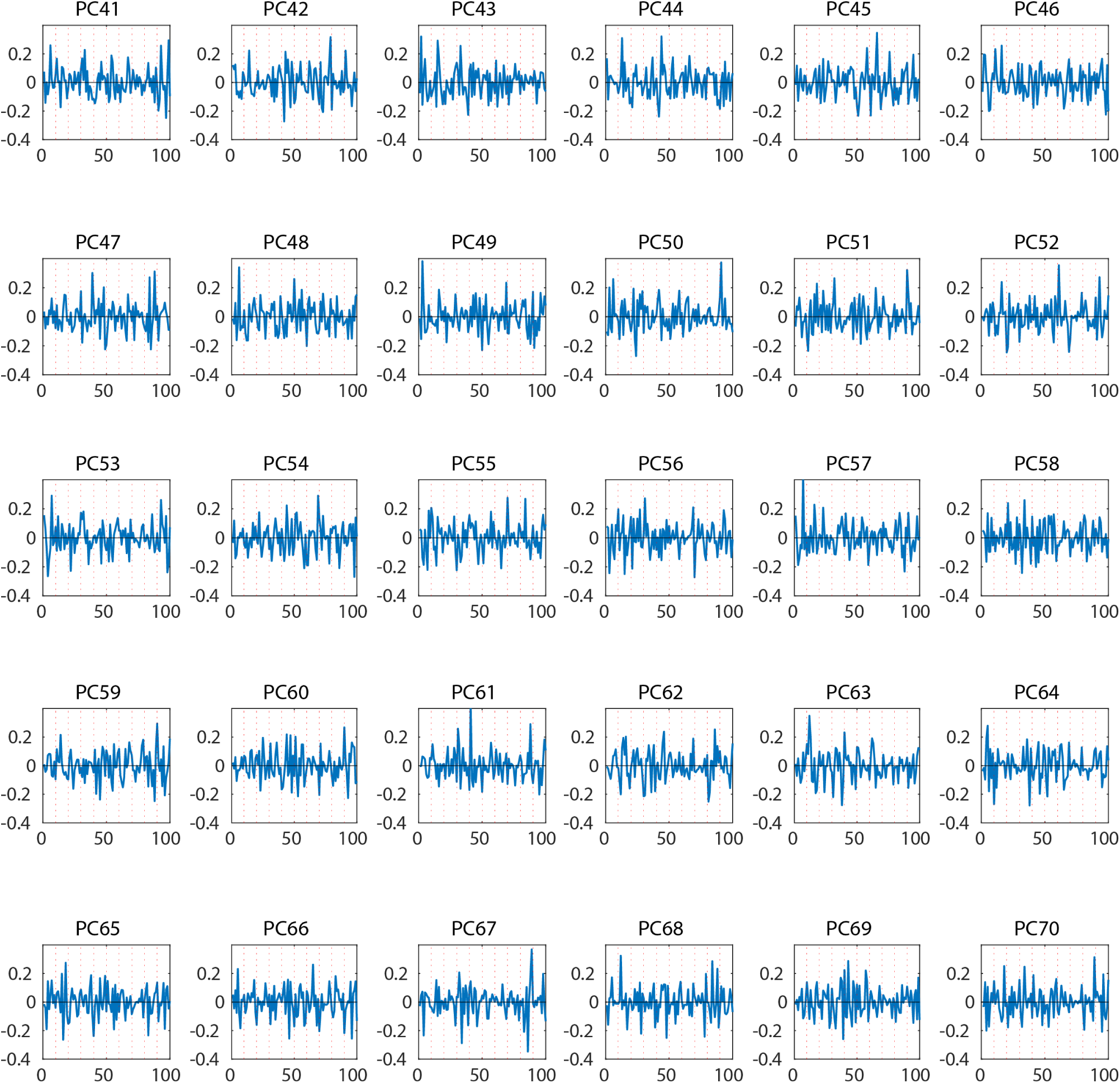
Coefficients for Principal Components 41 to 70 in MSC 100 node eFC. Red-dotted lines used for separating subjects’ scans.

**FIG. S8.**
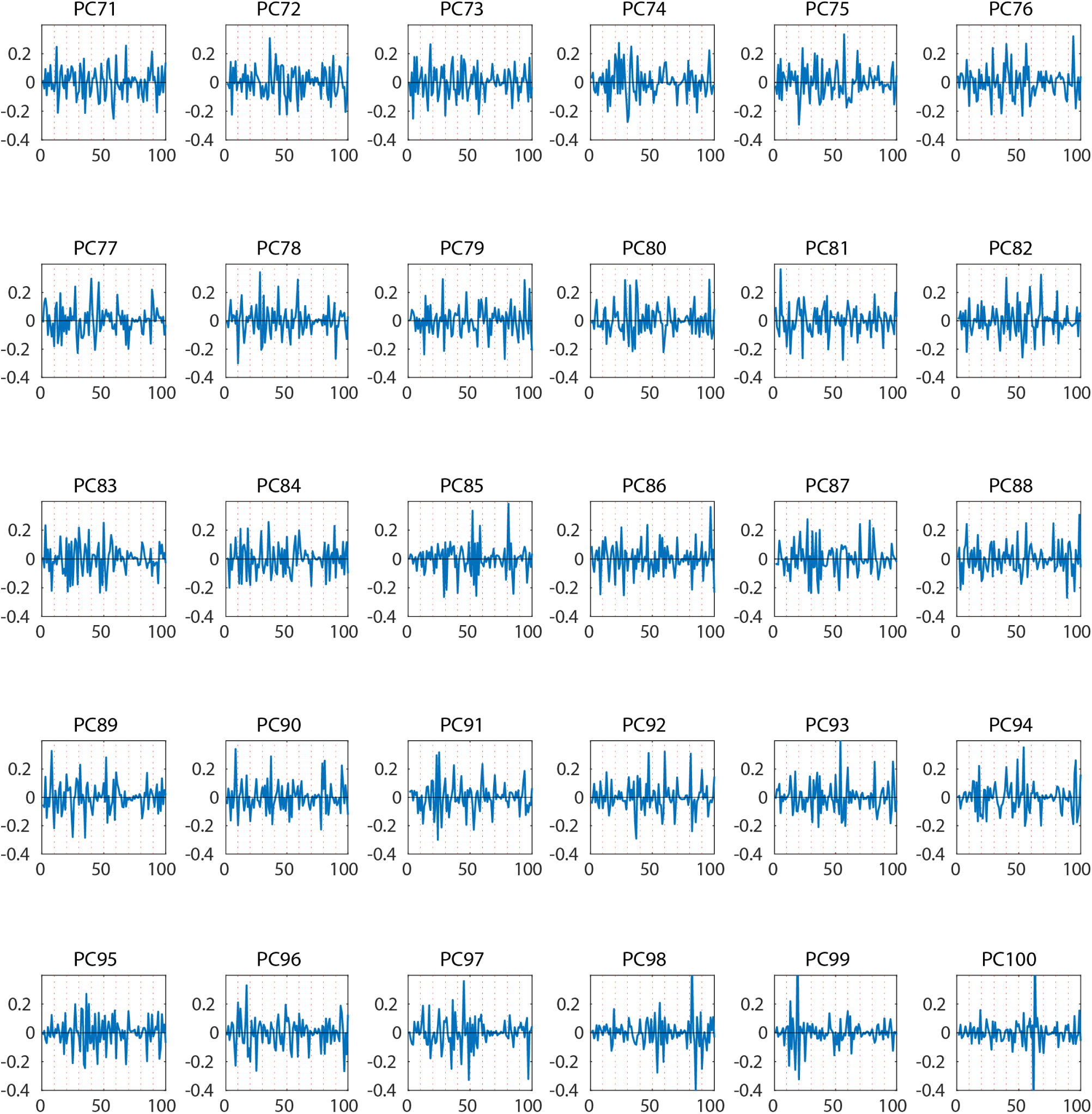
Coefficients for Principal Components 71 to 100 in MSC 100 node eFC. Red-dotted lines used for separating subjects’ scans.

